# Phosphorimidate Derivatives of Acyclovir; Antiviral Activity Against Canine Parvovirus *In Vitro*

**DOI:** 10.1101/598177

**Authors:** Payal M Oak, Akash S. Mali, Atul R Chopade, Govind Shukla

## Abstract

Canine parvovirosis is a very transmissible, severe and often deadly infectious disease of dogs caused by Type II canine parvovirus (CPV-2). Recently shows that Parvoviruses are very resistant to Acyclovir which is often used for HSV Chemotherapy in humans and various virucidal purposes in animal diseases in veterinary clinics and animal housing facilities. If acquiescence with vaccination programs and with appropriate decontamination plans is guaranteed, there should be no continuous, nor common, CPV-2 outbreaks. However, a continuous spread of CPV-2 infections is observed, even in shelters where an appropriate vaccination program is applied, and this is reason to provide an antiviral drug therapy. The aim of the present study was to Development of antiviral drugs with determination the effect of concentration of new chemical entity and analyze Acyclovir analogous against CPV-2 strains. A sensitive in vitro assay capable of measuring the infectivity of CPV-2 was employed to determine the efficacy of three different concentrations of 9-(2-hydroxyethomethyl)guanine phosphoromonomorpholidate. We successfully show that new compounds inhibit CPV-2 replication with exhibiting 50% inhibitory concentrations (IC_50_s) in the low-micromolar range (50μM).

## Introduction

Canine parvovirosis (CPV-2) is a very spreadable, severe and frequently mortal infectious disease which occurs in both domestic and wild dogs [1]. CPV-2, the etiological agent of canine parvovirosis, goes to the Parvoviridae family and Parvovirinae subfamily and it is included in Carnivore protoparvovirus 1 species together with Raccons parvovirus,Feline parvovirus and Mink enteritis virus [2]. CPV-2 is a small around 25 nm diameter, non-enveloped virus Composed by three major proteins surrounding a single-stranded linear DNA genome [3]. Usually, CPV-2 infects 2–12 weeks mature pups, particularly during the failure of maternally derived antibodies (MDA) [4]. Commonly, adults are resistant to CPV-2 infection due to reduced vulnerability or presence of immunity encouraged either by vaccination or previous infections [3]. The transmission route is the oronasal through direct or indirect contact with the faeces of infected dogs or contaminated fomites; indirect contact is facilitated by the environmental resistance of the virus. Diagnosis is conducted by real-time PCR,and NGS now days via detection of the CPV-2 DNA in faeces of infected pups. The virus is inactivated 60 seconds at 100°C while resists up to 7 h at 80 °C and 72 h at 56°C, in addition, CPV-2 resists for 14-21 days at 37°C and sometimes 6 months at room temperature [5]. Recent report shows that CPV-2 is resistant to most disinfectants while is sensitive to oxidizing agents, formalin, hydroxylamine, halogens, β-propiolactone [6]. Among with above compounds Acyclovir is showing effect against CPV infection in puppies. Acyclovir is guanine analogue frequently used antiviral Chemotherapy of low Cytotoxicity and mainly used for treatment of herpes simplex virus (HSV) infection (7,8). The purposes of this study were to record the therapeutically evaluation of an Acyclovir as a prophylactic Pharmacological agent for canine parvovirus. The aim of the present study was to determine the effect acyclovir analogous with different concentration and time frame against several CPV-2 strains *in vitro*.

## Material and Methods

### Drug and Cells

ACV(9-[(2-Hydroxyethoxy)methyl]guanine) was obtained as a gift sample from Merck, India. A72 adherent canine tumour cell line was obtained from American Type Culture Collection (Manassas, VA, USA). A72 cells were maintained in high glucose Dulbecco’s Modified Eagle’s Medium (Sigma Aldrich) supplemented by: 10% Fetal Calf Serum (Sigma Aldrich) and 1% L-Glutamine (Sigma Aldrich), 1% Penicillin-Streptomycin (Sigma Aldrich). A72 cell line was incubated at 37°C in humidified atmosphere with 5% CO2.

### Viruses

CPV-2 strains were used in the study: strain 2a 192/98 [11]. The viruses were spread on A-72 cells to obtain stock viruses for the subsequent experiments. Each stock virus was titrated on A- 72 cells. Briefly, after an incubation period of 48 hrs at 37°C, the infected cells were fixed with cold acetone and tested using CPV-2-specific canine antibodies and rabbit anti-dog IgG conjugated with fluorescein isothiocyanate (Sigma Aldrich). Viral titre, determined on A-72 cells, was 10^5.5^ tissue culture infectious doses TCID_50_/ml for strain 2a, 10^6^ TCID_50_/ml for strain 2b.

### Phosphorimidate Darivatives of Acyclovir

Phosphorus oxychloride (1.50 mol/equiv) was added to a suspension of ACV (1.00 mol/equiv) in triethylphosphate (1 ml) precooled to 18^°^C. The mixture was kept at 18^°^C for 4 h. Then, it was treated with the amidating agent and N,N-diisopropylethylamine (6.00 mol/equiv) in 1% aqueous dioxane (1 ml)). The reaction was maintained at +4^°^C for 2 h, neutralized by a saturated NaHCO3 solution (10 ml) pre-cooled to +4^°^C, and extracted with ether (10 ml). The aqueous extract was applied onto a DEAE-Toyopearl column and eluted with a linear gradient of NH_4_HCO_3_. The target fraction was concentrated by evaporation under vacuum; the residue was diluted with water, re-evaporated and additionally chromatographed on a RP-18 column and eluted with a linear gradient of acetonitrile in 0.05 M NH_4_HCO_3_. The fraction containing the target product was freeze-dried from water. it was diluted with sterile-distilled water to the final concentration[5]

### Toxicity test/Cell Proliferation assay

Using the MTS One Solution Cell Proliferation Assay, the metabolic activity of the cells can be used to determine the cytotoxicity of various substances. MTS [3- (4,5-dimethyldiazol-2-yl) −5- (3-carboxymethoxyphenyl) 2- (4-sulfophenyl) −2H-tetrazolium] is a chemical substance which is converted by protonation into a colored cell culture medium Substance (formazan). Metabolically active cells are responsible for the metabolism of NADPH or NADH protons, which can be absorbed by MTS. The amount of formazan is directly proportional to the number of living cells in a 96 “well” plate and can be determined by the absorbance at 490 nm. [11]

### Plaques Reduction assays

The virucidal activity was measured by in vitro incubation of viruses with the 9-(2- hydroxyethomethyl)guanine phosphoromonomorpholidate (ACV PMMPD) Briefly, 10^6^ p.f.u. of CPV strains were incubated for 30 min at RT or at 37°C with EC_50_ of 9-(2- hydroxyethomethyl)guanine phosphoromonomorpholidate (10,20,30….300 μm/ml), respectively. Simultaneously, the same amount of virus was incubated with ACV without Compound as control. The residual infectious viruses were quantified by viral plaque assays.[11]

## Data analysis

Three independent experiments were performed with each strain of CPV-2 and one representative set of data was shown for each strain. Student’s t test was used to evaluate statistical differences which were considered significant when P was <0.05. The EC50 values were calculated using GraphPad Prism software v. 5.01.

## Results

### Cytotoxicity assay

To assess a toxic effect of the substance on the cell lines used, the MTS solution was used. Using this tetrazolium salt, the metabolic activity could be measured by measuring the absorption at 490 nm and thus the vitality of the cells was concluded. For this test, A-72 cells were seeded in a 96 “well” plate with 2×104 cells in 100μl cell culture medium and the substances to be tested in the final concentrations of 30μM, 10μM, 3μM and 1μM in triplets on the cells. The substances dissolved in DMSO were diluted 1: 100 in the cell culture medium from the stock solutions (300nM, 100nM). The same volume (1μl) of DMSO was used as the control. The cells were cultured for four days with the substances in the Incubator at 37°C. and 5% CO2. After four days, 20μl of MTS solution per “well” were added to the cells and the cells were incubated with the MTS solution for 2 h in the Incubator. Subsequently, the absorbance at 490 nm was measured.

For the calculation of the EC50, the data of the titration from one approach of the triplets were used. Thus, three independent EC50 values per substance were obtained (Table no.1), which were subsequently averaged. The standard deviation was specified as an error. The EC50 of ACVPMMPD are in the micro molar range having the smallest mean effective concentration with 18.86μM CPV titer on incubation of infected cells with the active pharmacological substances. As a control, the virus titer was applied from cells treated with ACV.

**Table No.1.** Comparison of the calculated EC50 and IC50 values of the tested substances

| Compound | IC<sub>50</sub> | EC<sub>50</sub> |
| --- | --- | --- |
| Standard/ACV | 9.63±0.01 | 7.1±0.6 |
| ACVPMMPD |  |  |
|  | 17.41±0.65 | 18.86±0.1 |

### Viral Inhibition

In this first test ACV derivatives were identified as inhibitors of CPV replication. The substance which drowned the virus titer were tested in a further infection test in lower concentrations than in the first experiment to be able to calculate the mean effective concentration (EC_50_). The substances were expressed in ascending concentrations (0.1μM, 1μM, 10μM, 12μM, 14 μM,16 μM,18 μM….24 μM) The cells were then infected with CPV at a MOI of 0.05-0.1 and cultured for 3 days in the breeder compartment. These approaches were each executed in triplets. After the titration and subsequent cultivation of the cells for another three days and the infection herd was counted.

## Discussion and Conclusion

The advance of antiviral drugs is still in its beginning with rapid changes and progressive milestones come across almost daily. The last 30 years have been the most dynamic in the history of viral infections and their management. Unfortunately, antiviral drugs have been active for only a few groups of viruses up until now [8,9] Maximum antiviral drugs do not provide a cure, but rather allow control of the contamination. However, the limitations of Chemo therapy, with the high costs of drugs, make the need for anticipation even more urgent. Concerning to the effect of Acyclovir on treatment of CPV2 in experimentally diseased puppies. It was successes in preventing of CPV2 replication in puppies as showed in Fig. 2 and virus recovering, which revealed absences of viral particles in fecal swabs, leukopenia, lymphopenia and hypoproteinemia in compared to 2nd group and this supported by Piret et al., who mentioned that the Acyclovir diverges from earlier nucleoside analogues in covering only a partial nucleoside structure: the sugar ring is replaced with an open-chain structure[12,16] It is selectively converted into 9-(2-hydroxyethomethyl)guanine phosphoromonomorpholidate which is far more effective in phosphorylation than cellular thymidine kinase. Subsequently, the monophosphate form is further phosphorylated into the active triphosphate form, 9-(2-hydroxyethomethyl guanine phosphoromonomorpholidate, by cellular kinases. It is incorporated into viral DNA, resulting in chain termination. It has also been shown that viral enzymes cannot remove 9-(2- hydroxyethomethyl) guanine phosphoromonomorpholidate from the chain, which results in inhibition of further activity of DNA polymerase. Similar results were obtained by Gertrude who tested the antiviral effect of Acyclovir against herpes virus type-1 (which is a DNA virus as CPV) and these results come in complete agreement with obtained using Acyclovir against HV-1 in skin samples from experimentally infected mice by Piret et al. Detectable antiviral effect against canine virus and it was valuable to reduce the severity of CPV-2 *in vitro*

**Figure No.1.**
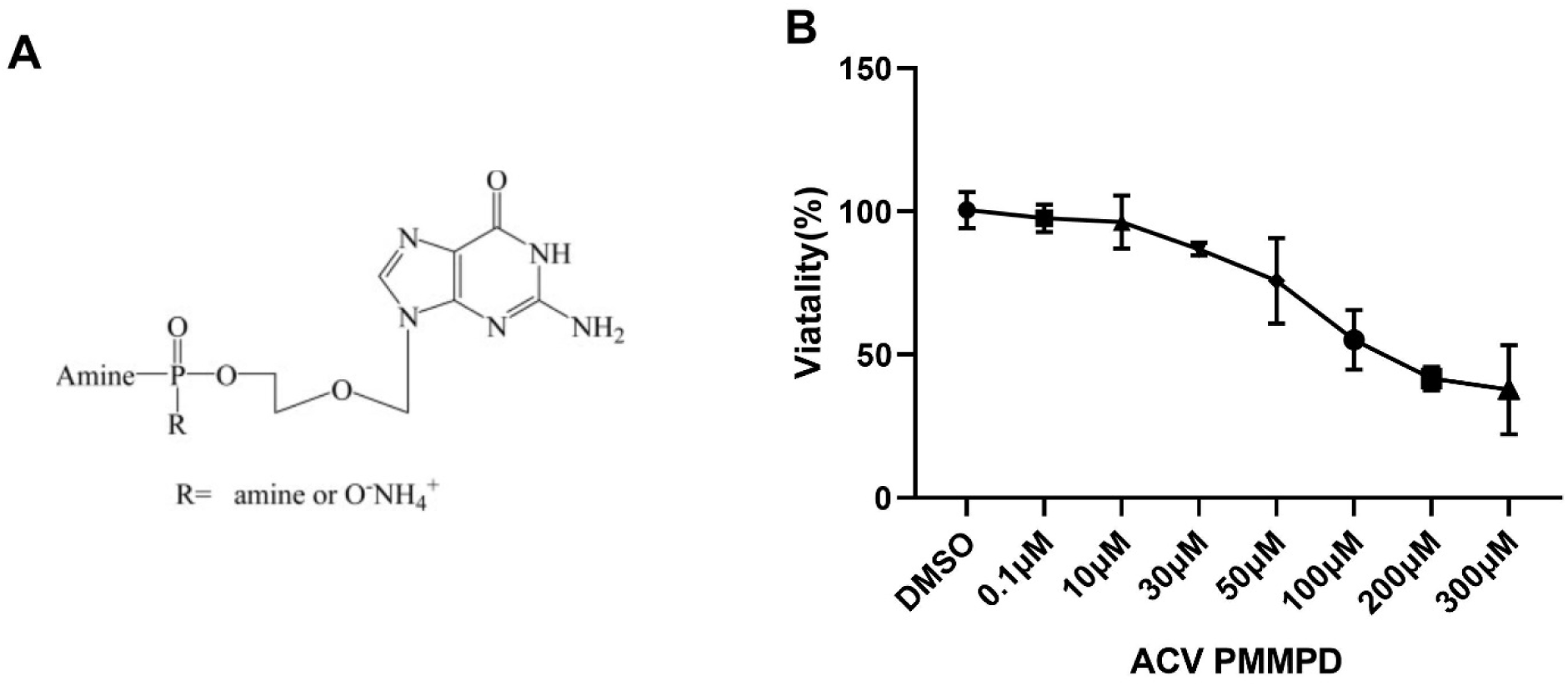
(A) New analogue 9-(2-hydroxyethomethyl)guanine phosphoromonomorpholidate (ACV PMMPD) Synthesized from Acyclovir, (B) Cytotoxic Assay, A-72 cells treated with various concentration(0.1,10,30,50,100,200,300 micromolar/ml) and assay carried out by using MTS solution, Absorbance values of A72 cell line culture after 48 hrs A72 cells (2×10^4^ cells/well) were sown and cultured at standard conditions (5% CO_2_, 37°C and humidified atmosphere).

**Figure No. 2.**
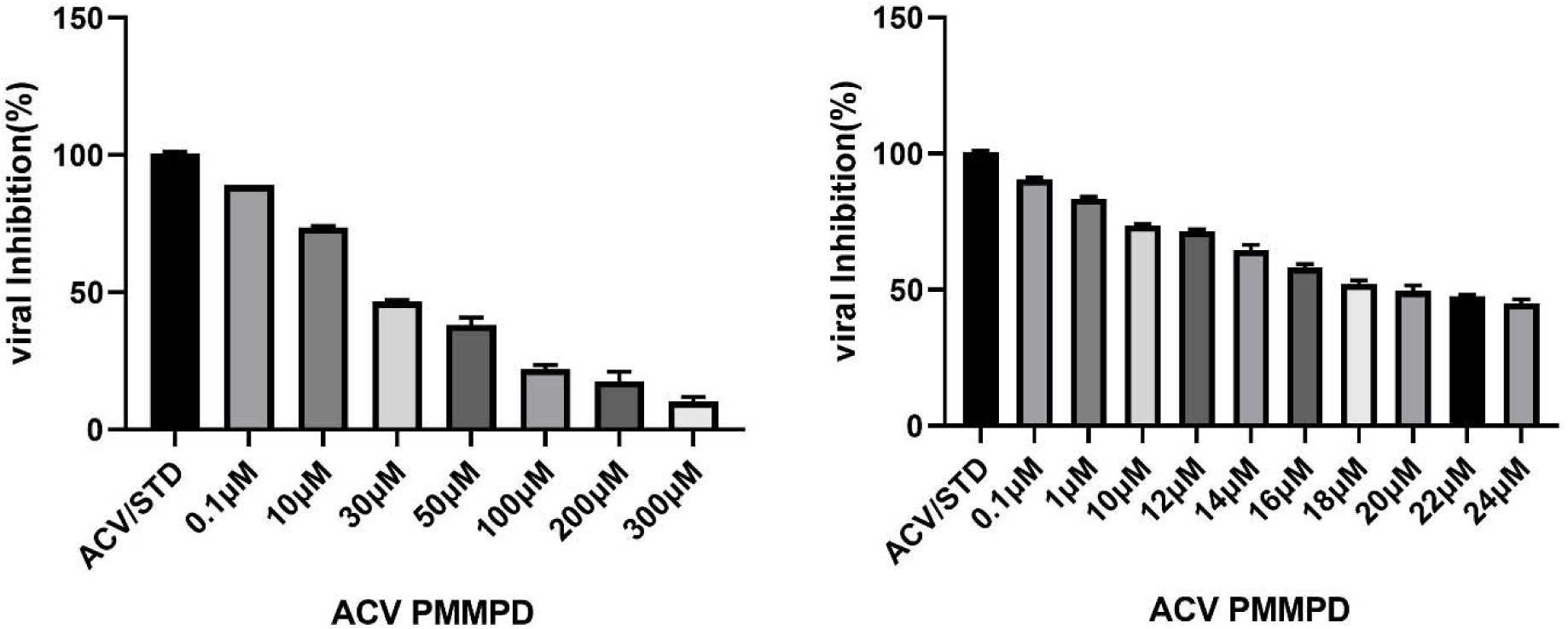
Following treatment after 24 Hrs, Graphical representation of the CPV strain titre on incubation of infected cells with the corresponding Concentration of ACV PMMPD. As a control, the virus titer was applied from cells treated with ACV. The standard deviation was reported as an error.

## Acknowledgments

The authors are deeply grateful to Professor Dr. Atul Chopade for encourage our team and Govind Shukla for continuous support.

## Reference

1. Appel MJ, Scott FW and Carmichael LE (1979) Isolation and immunization studies of a canine parvo-like virus from dogs with haemorrhagic enteritis. The Veterinary Record 105, 156–159.

2. Buonavoglia D et al. (2000) Antigenic analysis of canine parvovirus strains isolated in Italy. The New Microbiologica 23, 93–96.

3. Buonavoglia C et al. (2001) Evidence for evolution of canine parvovirus type 2 in Italy. Journal of General Virology 82, 3021–3025

4. Cavalli A et al. (2008) Evaluation of the antigenic relationships among canine parvovirus 2 variants. Clinical and Vaccine Immunology 15, 534–539.

5. Lisco, A.; Vanpouille, C.; Tchesnokov, E. P.; Grivel, J.-C.; Biancotto, A.; Brichacek, B.; Elliott, J.; Fromentin, E.; Shattock, R.; Anton, P.; Gorelick, R.; Balzarini, J.; McGuigan, C.; Derudas, M.; Gotte, M.; Schinazi, R. F.; Margolis, L. Cell Host Microbe 2008, 4, 260.

6. Decaro N and Buonavoglia C (2017) Canine parvovirus post-vaccination shedding: interference with diagnostic assays and correlation with host immune status. Veterinary Journal 221, 23–24.

7. Decaro N and Buonavoglia C (2012) Canine parvovirus-a review of epidemiological and diagnostic aspects with emphasis on type 2c. Veterinary Microbiology 155, 1–12.

8. Decaro N et al. (2005a) Clinical and virological findings in pups naturallyinfected by canine parvovirus type 2 Glu-426 mutant. Journal of Veterinary Diagnostic Investigation 17, 133–138

9. Greene CE and Decaro N (2012) Canine viral enteritis. In Greene CE (ed.), Infectious Diseases of the Dog and Cat, 4th Edn. St Louis, MO: Elsevier Saunders, pp. 67–80.

10. Lanave G et al. (2017) Ribavirin and boceprevir are able to reduce canine distemper virus growth in vitro. Journal of Virological Methods 248, 207–211.

11. McGavin D (1987) Inactivation of canine parvovirus by disinfectants and heat. Journal of Small Animal Practice 28, 523–535.

12. Michaela Rumlová, Tomáš Ruml, In vitro methods for testing antiviral drugs, Biotechnology Advances, Volume 36, 3, 2018, 557–576

13. Scott FW (1980) Virucidal disinfectants and feline viruses. American Journal of Veterinary Research 41, 410–414.

14. Spibey N, Greenwood NM, Sutton D, et al. Canine parvovirus type 2 vaccine protects against virulent challenge with type C virus. Veterinary Microbiology. 2008;128:48□55.

15. Oie S et al. (2011) Disinfection methods for spores of Bacillus atrophaeus, B. anthracis, Clostridium tetani, C. botulinum and C. difficile. Biological & Pharmaceutical Bulletin 34, 1325–1329.

16. Otto CM, Drobatz KJ, Soter C. Endotoxemia and tumor necrosis factor activity in dogs with naturally occurring parvoviral enteritis. J Vet Intern Med. 1997;11(2):65□70

17. Piret J, Desormeaux A, Gourde P, et al. Efficacies of topical formulation of foscarnet and acyclovir ointment (Zovirax) in a murine model of cutaneous herpes simplex virus type 1 infection. Antimicrob Agents Chemother. 2000;44(1):30□38.

